# The impact of systemic blockade of dopamine receptors on the acquisition of two-way active avoidance in rats

**DOI:** 10.1101/2024.11.29.626069

**Authors:** L. Vercammen, A. Lopez-Moraga, T. Beckers, B. Vervliet, L. Luyten

## Abstract

Active threat avoidance is a core aspect of adaptive and maladaptive behavior, yet its underlying mechanisms are not fully understood. Prior studies concluded that pharmacologically blocking dopaminergic receptors (DRs) disrupted avoidance acquisition, but it remains unclear whether such effects on learning persist during a drug-free follow-up test. To assess the involvement of D1R and D2R in avoidance acquisition, we conducted two experiments. In Experiment 1, thirty-six male Wistar rats underwent a single avoidance training session involving 30 tone-shock pairings. Rats could avoid the shock by moving to the opposite compartment of the shuttle box. Twenty minutes before training, rats received either D1R antagonist SCH 23390 (0.05 mg/kg), D2R antagonist sulpiride (20 mg/kg), or vehicle. While sulpiride did not affect avoidance, 0.05 mg/kg SCH 23390 significantly reduced the number of avoidance responses. In a separate test, 0.05 mg/kg SCH 23390 also reduced locomotor activity. In Experiment 2 (N = 24), a lower dose of SCH 23390 (0.025 mg/kg) was administered, and a drug-free avoidance test under continued reinforcement was added 24 hours later to test for sustained effects of D1R blockade on avoidance in the absence of acute drug effects. Although animals avoided less with SCH 23390 in the system, this effect did not persist 24 hours later, suggesting that effects of D1R blockade during avoidance training might reflect an acute disruption of secondary processes involved in the performance of avoidance behavior rather than an actual impairment of avoidance learning.

## 1. Introduction

Active avoidance of threat is a hallmark of adaptive fear, e.g., most persons will try to avoid a stray dog that is aggressively barking. But avoidance can lose its adaptive function when it is not calibrated to the actual level of threat (Krypotos et al., 2015). When the barking dog is safely behind a fence, it is not necessary to take a detour. Excessive avoidance is a key characteristic of various mental health disorders, including anxiety disorders, obsessive-compulsive disorder and depression (Bardeen et al., 2014; Gillan et al., 2014; Ottenbreit et al., 2014). Given the role of avoidance in pathological functioning, elucidating the neurotransmitter systems involved in how individuals learn to avoid may help to understand the pathophysiology of anxiety disorders and related psychiatric conditions (Krypotos et al., 2015).

One potential target for this is the mesolimbic dopaminergic system. Dopamine has been implicated in learning, and more specifically, in linking environmental cues with outcomes that are better than expected, thereby guiding behavior to respond appropriately to those cues (Schultz, 2007). During avoidance learning, animals learn to link the performance of a behavioral response with the omission of an aversive outcome that would otherwise occur (Ledoux et al., 2017). For example, in a two-way active avoidance task (2WAA), rats learn that they can move to the opposite compartment of a shuttle box during the presentation of a warning signal (e.g., a tone) in order to prevent an upcoming aversive stimulus (e.g., a foot shock). This process is thought to be dopamine-dependent, given that the absence of the aversive event can be classified as an outcome that was “better than expected”, resulting in a reward prediction error (rPE; Kim et al., 2006). Such rPE is thought to reinforce the instrumental response (Schutz, 2007), thereby resulting in the acquisition of the avoidance behavior.

Prior research supports a role for the dopamine system in avoidance learning. Dopamine release, as measured via in-vivo microdialysis in the nucleus accumbens, follows a typical rPE pattern, with more release during early avoidance trials when the absence of shock is surprising and less release during later avoidance when the omissions have become expected (Dombrowski et al., 2013). Other studies have demonstrated a role for dopamine by blocking dopaminergic receptors during avoidance acquisition, through pharmacological manipulations. Several studies have observed impaired avoidance acquisition after pretraining systemic administration of D1R (e.g., SCH 23390; Wietzikoski et al., 2012; Reis et al., 2004) or D2R antagonists (e.g., sulpiride; Boschen et al., 2011; Reis et al., 2004), in line with the hypothesis that avoidance acquisition requires dopamine signaling. Together, these studies suggest that absence of the aversive stimulus during early avoidance trials triggers the release of dopamine in the nucleus accumbens, which then binds to specific dopamine receptors to support avoidance learning.

However, it is currently unclear whether the effects of dopamine receptor blockade persist during drug-free follow-up tests under continued reinforcement. One study (focusing on D1R, 9-10 animals per group) indeed found a persistent impairment during a drug-free follow-up test 24 hours later (Wietzikoski et al., 2012), but another study (focusing on D2R, 8-10 animals per group) did not observe a lasting effect on avoidance performance (Boschen et al., 2011). The majority of studies did not include a drug-free follow-up test at all (Reis et al., 2004; Carvalho et al., 2009; Diaz-Veliz et al., 1998). This raises the question whether dopamine receptor blockade disrupts the instrumental learning process per se, or rather affects secondary processes like motivation (Bromberg-Martin et al., 2010; Berke, 2018) or locomotor activity (Meyer et al., 1993) that can also impact the performance of avoidance behavior. If DR blockade indeed disrupts the instrumental learning process that is necessary for the acquisition of the avoidance response, rats treated with a dopamine antagonist should still display less avoidance than a vehicle control early on during a drug-free follow-up test, which is not consistently observed. Note that the observation of a sustained effect on a drug-free test does not rule out that effects of DR blockade could be mediated by impaired motivation or locomotion, as such impairments, through their effects on the execution of avoidance behavior, could also prevent proper avoidance acquisition. Conversely, however, if effects of DR blockade are not sustained on a drug-free test, this would suggest that such effects do not reflect an impairment in avoidance learning but a pure performance effect due to disrupted motivation or motor behavior only.

The inconsistency between studies could be related to the sensitivity of the analyses that were used. When collapsing avoidance responses across an entire test session, which usually consists of 20 to 50 trials, transient effects during early trials could be missed. Importantly, if administration of a dopamine antagonist prior to training indeed impairs the acquisition of avoidance, one would expect drug-treated rats to show fewer avoidance responses than vehicle controls particularly during the initial stages of the drug-free test, catching up during later stages of the test.

Through a combination of pharmacological manipulations and behavioral assessments, the present study was set up to test whether systemic administration of a DR antagonist before avoidance training impairs the acquisition of avoidance in a two-way active avoidance task. Experiment 1 served as a replication study to ascertain whether D1R and D2R blockade impair avoidance when administered before training (Reis et al., 2004, Boschen et al., 2011; Wietzikoski et al., 2012). Rats received systemic administrations of either D1R antagonist SCH 23390 (0.05 mg/kg), D2R antagonist sulpiride (20 mg/kg) or vehicle (saline + 17% DMSO) and the effect on avoidance acquisition was evaluated. In Experiment 2, a drug-free test was added 24 hours after acquisition to assess whether the deficit caused by D1R blockade was persistent, in particular during the early trials of the session, as a critical test of the notion that this deficit reflects an impairment in instrumental learning.

## 2. Methods and Materials

### 2.1. Preregistration and data availability

The experiments were preregistered on the Open Science Framework (https://osf.io/wypab/). The raw data files, processed data files, video files and R scripts generated for the current study are available via the same link.

### 2.2. Subjects

Sixty male Wistar rats (weighing between 275 – 300 grams upon arrival in the lab) were purchased from Janvier Labs (Le Genest Saint Isle, France). Rats were housed per two in standard animal cages with a red cylinder as cage enrichment and kept under conventional laboratory conditions (12-h day-night cycle; lights on 7:00-19:00; 22°C) with ad libitum access to food and water. The number of animals per group was calculated by a power analysis using effect sizes found in the literature (*d* = 1.35, adjusted to 1.20) to achieve a power of .80 (Reis et al., 2004; G-Power version 3.1). Experiments were conducted between 8:30 and 17:00. Rats were handled briefly on two consecutive days, prior to behavioral testing. All experiments were approved by the KU Leuven animal ethics committee (project P201/2020) and conducted in accordance with the Belgian and European laws, guidelines and policies for animal experimentation, housing and care (Belgian Royal Decree of 29 May 2013 and European Directive 2010/63/EU on the protection of animals used for scientific purposes of 20 October 2010).

### 2.3. Apparatus

#### 2.3.1. Two-way active avoidance apparatus

Two-way active avoidance training took place in two identical rectangular shuttle boxes (50.8 cm x 25.4 cm x 30.5 cm), each placed inside larger sound-attenuating cubicles (72.4 cm x 41.1 cm x 43.2 cm, Coulbourn Instruments, Pennsylvania, USA). The side walls and ceilings of the shuttle boxes were made of metal, whereas the front and back walls were clear Plexiglas. The shuttle box was divided into two equal compartments by a metal divider placed halfway along the length of the shuttle box (Coulbourn Instruments). Rats could freely move from one side of the box to the other via a passage (8 cm width) in the divider. The floor of the shuttle box was a stainless steel grid, through which scrambled foot shocks could be delivered (0.6 mA, 10 s) that served as the unconditioned stimulus (US). In each compartment of the shuttle box, a speaker was mounted to the side wall to deliver a 3-kHz tone (presented at +/-75 dB) that served as the conditioned stimulus (CS). A 0.5-W house light (white light) was mounted to the ceiling of each compartment and served as a safety signal. This signal was presented whenever the rat successfully avoided or escaped the foot shock. When the safety signal was not on, the shuttle box was unilluminated. Two photocell sensor bars, one in each side of the shuttle box, ensured automatic detection of the position of the rat throughout the experiment. All behavioral sessions were videorecorded via an infrared camera (HD IP camera, Foscam C1, Shenzhen, China) mounted to the ceiling of the isolation cubicle.

#### 2.3.2. General locomotor activity test

The general locomotor activity test took place in a dark grey arena (38 cm x 38 cm x 60 (H) cm). There was no visible light throughout the session, which was videorecorded via an infrared camera (HD IP camera, Foscam C1, Shenzhen, China) mounted above the test arena. The recorded videos were analyzed via Ethovision software XT (version 16, Noldus, Wageningen, the Netherlands).

#### 2.3.3 Accelerating rotarod

The accelerating rotarod (Ugo Basile, Gemonio, Italy) consisted of four sections (8.7 cm) to allow for simultaneous testing of four rats. Height of the rod (diameter: 6 cm) was 30 cm. During each trial, the rod accelerated from 5-40 rpm in 4 minutes (Luyten et al., 2017). Fall time was automatically recorded via the rotarod hardware (Ugo Basile).

### 2.4. Drug administration

Rats were intraperitoneally injected with either D1R antagonist SCH 23390 (0.05 mg/kg or 0.025 mg/kg, Tocris Bioscience), D2R antagonist sulpiride (20 mg/kg, Sigma Aldrich) or vehicle (0.9% physiological saline + 17% DMSO). According to a literature review, SCH 23390 is most commonly administered intraperitoneally at doses ranging from 0.0625 to 0.1 mg/kg (Wietzikoski et al., 2012; Reis et al., 2004; Diaz-Veliz et al., 1998), with the 0.05 mg/kg dose producing reliable behavioral effects. Sulpiride has most commonly been administered at doses ranging from 10 to 40 mg/kg (Boschen et al., 2011; Diaz-Veliz, 1998; Santacana et al., 1975; Reis et al., 2004; Carvalho et al., 2009), with 20 mg/kg producing reliable behavioral effects. All drugs and vehicle were administered at a volume of 4 ml/kg. Rats were randomly assigned to a group and injected 20 minutes prior to the start of the avoidance training session. The experimenter was blinded to the drug condition.

### 2.5. Behavioral procedures

For an overview of the experimental procedures, see Figure 1.

**Figure 1.**
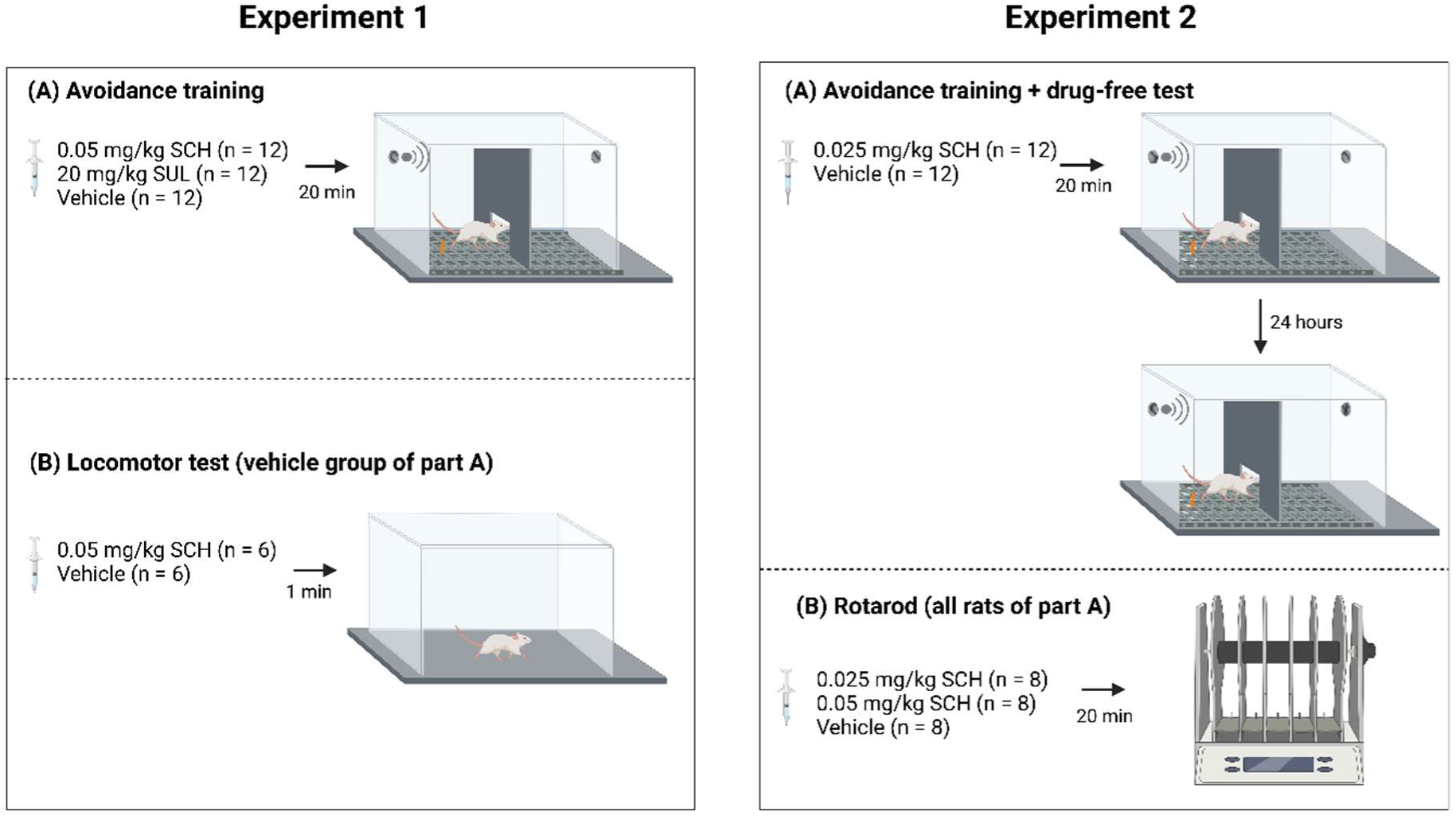
Overview of the behavioral procedures of Experiment 1 and Experiment 2. SCH = D1R antagonist SCH 23390, SUL = D2R antagonist sulpiride, vehicle = 0.9% physiological saline + 17% DMSO.

#### 2.5.1. Experiment 1

##### 2.5.1.1. Two-way active avoidance training

Twenty-four hours before the start of avoidance training, all rats were habituated to the shuttle box for 1 hour. On the following day, the rats were subjected to a single avoidance training session.

Twenty minutes prior to the start of avoidance training, the rats were administered with either the vehicle (n = 12), a D1R antagonist (0.05 mg/kg SCH 23390, n = 12) or a D2R antagonist (20 mg/kg sulpiride, n = 12). The avoidance training session started with a 5-minute acclimation period during which no stimuli were delivered. After acclimation, the rats were given 30 presentations of the CS and US, separated by an intertrial interval (ITI) averaging around 1 minute (30 s – 90 s). The CS lasted a maximum of 20 seconds and was followed immediately by the US which lasted a maximum of 10 seconds. If the rat shuttled to the other compartment during CS presentation, the CS was terminated and no shock was given, i.e., the US was avoided and the trial was labeled as an avoidance response. If the rat crossed to the other compartment during the presentation of the US, the shock was terminated and the trial was labeled as an escape response. After each successful avoidance or escape response, rats were presented with a safety signal (house light for 5 seconds). The total duration of the training session was 45 minutes. The number of avoidance responses, escape responses, escape failures (i.e., the rat remained in the compartment throughout the US presentation) and intertrial crossings (i.e., crossings during the ITIs) were recorded via Graphic State 4 software (Coulbourn Instruments). The delivery of the stimuli and data collection was fully automated by a PC running Graphic State 4 software (Coulbourn Instruments).

##### 2.5.1.2. General locomotor activity test

Seven days after avoidance acquisition, the vehicle rats of Experiment 1 were reused to evaluate the effect of 0.05 mg/kg SCH on locomotor activity. The rats were introduced to the arena for a 1-hour session and the total distance traveled (in cm) was recorded in five-minute bins. Rats were injected with either vehicle (0.9% physiological saline + 17% DMSO, n = 6) or 0.05 mg/kg SCH 23390 (n = 6), and immediately placed in the test arena.

#### 2.5.2. Experiment 2

##### 2.5.2.1. Two-way active avoidance training

Avoidance training proceeded as described above.

##### 2.5.2.2. Two-way active avoidance test

Twenty-four hours after avoidance training, the rats were subjected to an additional (drug-free) avoidance test under continued reinforcement. The experimental parameters were equal to those used during avoidance training.

##### 2.5.2.3. Accelerating rotarod

Seven days after the avoidance test, the rats were reused to evaluate the effect of 0.025 mg/kg and 0.05 mg/kg SCH 23390 on motor function using an accelerating rotarod training and test procedure. Each rat received a different dose than what they had previously received before avoidance acquisition. Rats first underwent three days of training (consisting of 3 consecutive trials on each day) on the accelerating rotarod, so that a stable rotarod performance was established. On the fourth day, the rats were administered with either 0.05 mg/kg SCH 23390 (n = 8), 0.025 mg/kg SCH 23390 (n = 8) or vehicle (0.9% physiological saline + 17% DMSO, n = 8), 20 minutes prior to an accelerating rotarod test (consisting of 3 consecutive trials). The latency to fall off the rotarod was automatically recorded.

### 2.6. Statistical analyses

All analyses were preregistered on the Open Science Framework, unless stated otherwise. Data are expressed as means ± SEM and analyzed using R statistical software (version 3.5.1). The mean number of avoidance responses was calculated for each session per group and analyzed using one-sided independent samples t-tests (assessing if drug < vehicle). The mean number of escape responses and escape failures was analyzed exploratively using independent two-sided t-tests. The mean number of avoidance responses was calculated per block of 10 trials within a session and analyzed using a mixed design ANOVA (repeated-measures factor: Block, between-subjects factor: Group). To further test our hypotheses, we conducted a non-preregistered analysis for which the avoidance responses were analyzed on a trial-by-trial basis by coding the data in binary format (avoidance = 1, escape/escape failure = 0) and applying a generalized linear mixed-effects model to the binary outcome with fixed-effects parameters Trial, Group and the interaction between Trial and Group, and a random effects parameter Subject to account for within-subject variability due to the repeated measures design, using the R-package lme4 (https://cran.r-project.org/package=lme4). Finally, the mean number of intertrial crossings was analyzed and compared between groups using two-sided independent samples t-tests. For the locomotor activity test and accelerating rotarod test, mixed design ANOVAs were applied (within-subjects factor = Time bin or Phase, between-subjects factor = Group). Preregistered secondary analyses can be found in the Supplementary Material. The average latency to avoid the US, as well as the average latency to escape the US were analyzed using two-sided independent samples t-tests (Suppl. Fig. 1). Moreover, the latency to avoid the US was calculated per block of 10 trials within a session and analyzed using a mixed design ANOVA (repeated-measures factor: Block, between-subjects factor: Group). The average number of seconds of freezing during CS1 and during each even CS presentation (CS2, CS4,…) was analyzed using a mixed design ANOVA (repeated-measures factor: CS, between-subjects factor: Group). Moreover, the average number of seconds of freezing across all even CS presentations was analyzed using two-sided independent samples t-tests (Suppl. Fig. 2). Finally, the average number of shuttles during the 5-minute habituation period of both the avoidance training session and the avoidance test session of Experiment 2 were analyzed using two-sided independent samples t-tests (Suppl. Fig. 3).

## 3. Results

### 3.1. Experiment 1

#### 3.1.1. Two-way active avoidance training

Rats were intraperitoneally injected with D1R antagonist SCH 23390 (0.05 mg/kg, n = 12), D2R antagonist sulpiride (20 mg/kg, n = 12), or vehicle (0.9% physiological saline + 17% DMSO, n = 12), 20 minutes prior to a single avoidance training session.

Rats given 0.05 mg/kg SCH 23390 (SCH) showed fewer avoidance responses compared to the control group (*t*(11.33) = -7.823, *p* < .001, *d* = 0.769; Figure 2A). In contrast, there was no significant effect of 20 mg/kg sulpiride (SUL) during avoidance acquisition (*t*(22) = 1.285, *p* = .106, *d* = 0.524; Figure 2A). Exploratively, potential differences in the escape responses and escape failures were also analyzed, due to the observation of a relatively high number of escape failures in several subjects. There was no significant difference in the average number of escape responses for both the SCH group (*t*(22) = -0.842, *p* = .409, *d* = 0.344) and the SUL group (*t*(22) = -1.335, *p* = .196, *d* = -0.545, Figure 2B), compared to the control group. There were significantly more escape failures following SCH administration (*t*(22) = 4.624, *p* < .001, *d* = 1.888), but not following SUL administration (*t*(22) = - 0.138, *p* = .891, *d* = 0.408; Figure 2C), compared to the control group.

**Figure 2.**
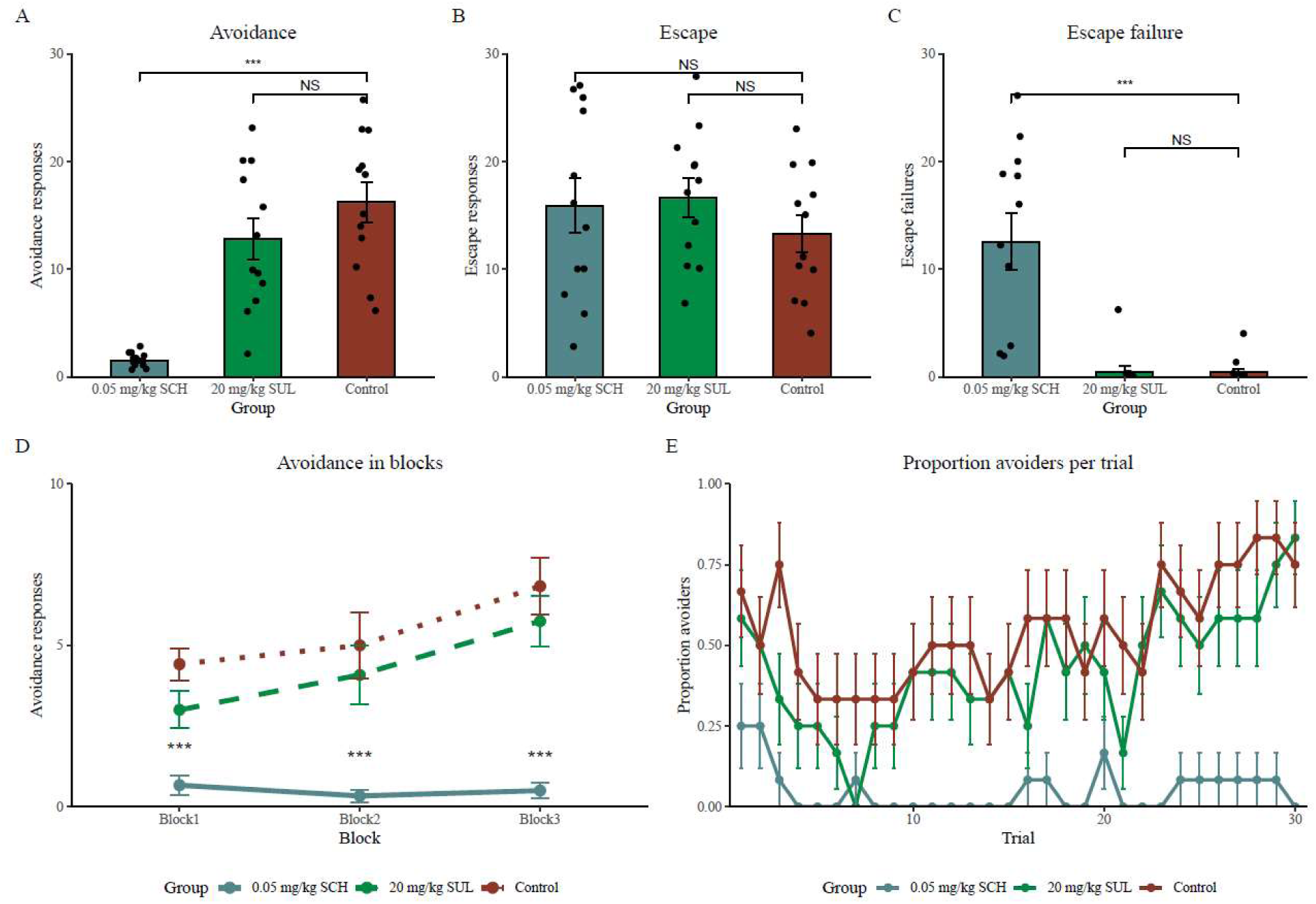
Experiment 1: Two-way active avoidance training results. (A) Mean (± SEM) number of avoidance responses, (B) escape responses and (C) escape failures for the 0.05 mg/kg SCH (n = 12), 20 mg/kg SUL (n = 12) and control (n = 12) groups. *** *p* < .001. (D) Mean (± SEM) number of avoidance responses in blocks of 10 trials for the 0.05 mg/kg SCH, 20 mg/kg SUL and control groups. *** *p* < .001. (E) Proportion of avoiders per trial ± SEM for the 0.05 mg/kg SCH, 20 mg/kg SUL and control groups. Note that the proportions of the first few trials are relatively high since rats did not yet experience the CS-US association, and mere exploration of the opposite compartment during the tone was also counted as an avoidance response.

When analyzing the data of the avoidance training session in three blocks of 10 trials each, we found a significant main effect of Group (*F*(2,33) = 25.089, *p* < .001, ƞ*_p_²* = 0.603), a significant main effect of Block (*F*(2,66) = 8.177, *p* < .001, ƞ*_p_²* = 0.199), and a trend towards a significant Group by Block interaction effect (*F*(4,66) = 2.383, *p* = .060, ƞ*_p_²* = 0.126; Figure 2D). Post-hoc testing showed significantly fewer avoidance responses in the SCH group compared to the control group, in all blocks (*ps* < .001), but no significant differences between the control and the SUL group.

Finally, we conducted an exploratory generalized linear mixed model (GLMM) analysis to examine the effects of Trial and Group (reference group: control group) on the binary avoidance data (avoidance = 1, escape/escape failure = 0), with a random intercept for each subject. The model was fit using a binomial distribution with a logit link function and optimized using the Laplace approximation. The logistic regression coefficients indicated a significant effect of Trial (*β* = 0.054, *SE* = 0.014, *Z* = 3.901, *p* < .001) and a significant effect for the SCH group (*β* = -2.164, *SE* = 0.616, *Z* = -3.512, *p* < .001) suggesting that, compared to the control group, the SCH group was significantly less likely to avoid. The effect of the SUL group was not significant (*β* = -0.754, *SE* = 0.484, *Z* = - 1.558, *p* = .120). Finally, we also observed a significant negative interaction between Trial and the SCH group (*β* = -0.076, *SE* = 0.031, *Z* = -2.465, *p* = .014), indicating that the effect of Trial on the outcome was attenuated in the SCH group. This interaction was not significant for the SUL group (*β* = 0.012, *SE* = 0.020, *Z* = 0.614, *p* = .539). Overall, these findings suggest a reduction in avoidance responses for rats that were given 0.05 mg/kg SCH 23390, but not for rats that were given 20 mg/kg sulpiride, compared to a vehicle group.

#### 3.1.2. General locomotor activity test

To evaluate whether there were any effects of 0.05 mg/kg SCH and 20 mg/kg sulpiride on locomotor activity, we first evaluated the number of intertrial crossings during the avoidance training session. The SCH group shuttled significantly less during the ITIs than the control group (*t*(22) = - 4.907, *p* < .001, *d* = -2.003, Figure 3A). There was no difference in ITI crossings between the SUL and control groups (*t*(22) = 0.620, *p* = .542, *d* = 0.253, Figure 3A). This reduction in locomotor activity during the ITIs following 0.05 mg/kg SCH administration potentially explains why rats of this group performed fewer avoidance responses, compared to vehicle administered rats. Intact locomotor abilities are a prerequisite to adequately acquire and perform the two-way active avoidance response, as rats need to shuttle in order to avoid the foot shock.

**Figure 3.**
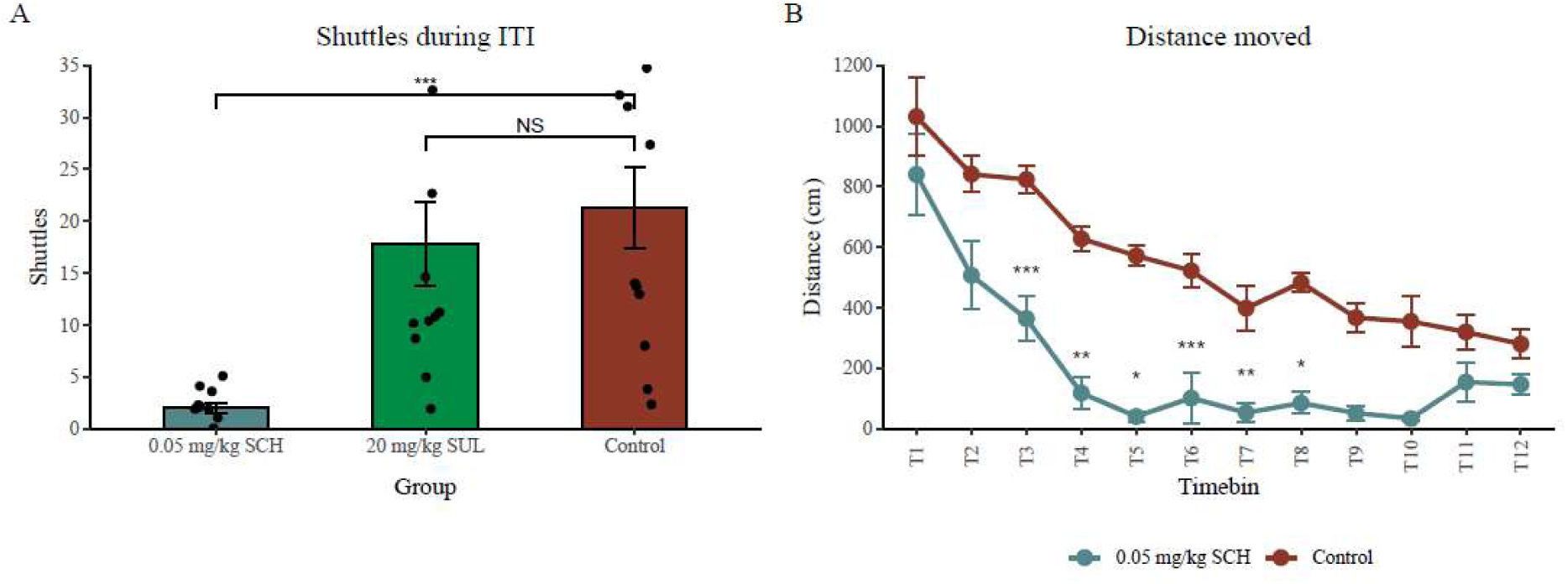
(A) Mean number of shuttles ± SEM during the ITIs of the avoidance training session for the 0.05 mg/kg SCH (n = 12), 20 mg/kg SUL (n = 12) and control (n = 12) groups. *** *p* < .001 (B) Mean distance moved (cm) during the general locomotor activity test ± SEM for the 0.05 mg/kg SCH group (n = 6) and the control group (n = 6) in time bins of 5 minutes each. *** *p* < .001, ** *p* < .01, * *p* < .05.

To further explore the reduced locomotion that was observed following 0.05 mg/kg SCH administration, a general locomotor activity test was performed. Rats received either 0.05 mg/kg SCH 23390 (n = 6) or vehicle (0.9% physiological saline + 17% DMSO, n = 6) immediately prior to being placed in the arena for 1 hour. The distance moved (cm) was analyzed in time bins of 5 minutes, and showed a significant main effect of Time bin (*F*(11,110) = 25.765, *p* < .001, ƞ*_p_²* = 0.720), suggesting that animals moved less towards the end of the session (Figure 3B). Additionally, there was a significant effect of Treatment (*F*(1,10) = 132.093, *p* < .001, ƞ*_p_²* = 0.930) and a significant Time bin by Treatment interaction effect (*F*(11,110) = 2.007, *p* = .034, ƞ*_p_²* = 0.167). Post-hoc testing showed significantly less distance moved for rats that received 0.05 mg/kg SCH 23390 in Time bins 3 – 8 (i.e., 10 to 40 min after injection), compared to rats that received the vehicle.

### 3.2. Experiment 2

#### 3.2.1. Two-way active avoidance training

Due to multiple escape failures observed following administration 0.05 mg/kg SCH 23390 in Experiment 1, as well as its clear suppression of locomotor activity, Experiment 2 aimed to evaluate the effect of a lower dose of SCH 23390 (0.025 mg/kg) on avoidance learning. This dose has previously been shown to acutely decrease the number of avoidance responses, but its long-term effects have not yet been studied (Reis et al., 2004; Diaz-Veliz et al., 1998). To evaluate whether SCH 23390 indeed impairs instrumental learning per se, rather than acutely affecting secondary processes, we not only investigated the effects during a first avoidance acquisition session, but also added a drug-free avoidance test under continued reinforcement 24 hours after training.

Twenty minutes prior to the avoidance training session, rats received either 0.025 mg/kg SCH (n = 12) or vehicle (saline + 17% DMSO, n = 12). Rats that received 0.025 mg/kg SCH showed significantly less avoidance responses, compared to the control group during avoidance training (*t*(22) = 2.054, *p* = .026, *d* = 0.838, Figure 4A). Administration of 0.025 mg/kg SCH did not result in a significant difference in escape responses (*t*(22) = -1.362, *p* = .187, *d* = 0.424; Figure 4B), nor in a significant increase in escape failures (*t*(22) = -1.901, *p* = .071, *d* = -0.776; Figure 4C), compared to the control group.

**Figure 4.**
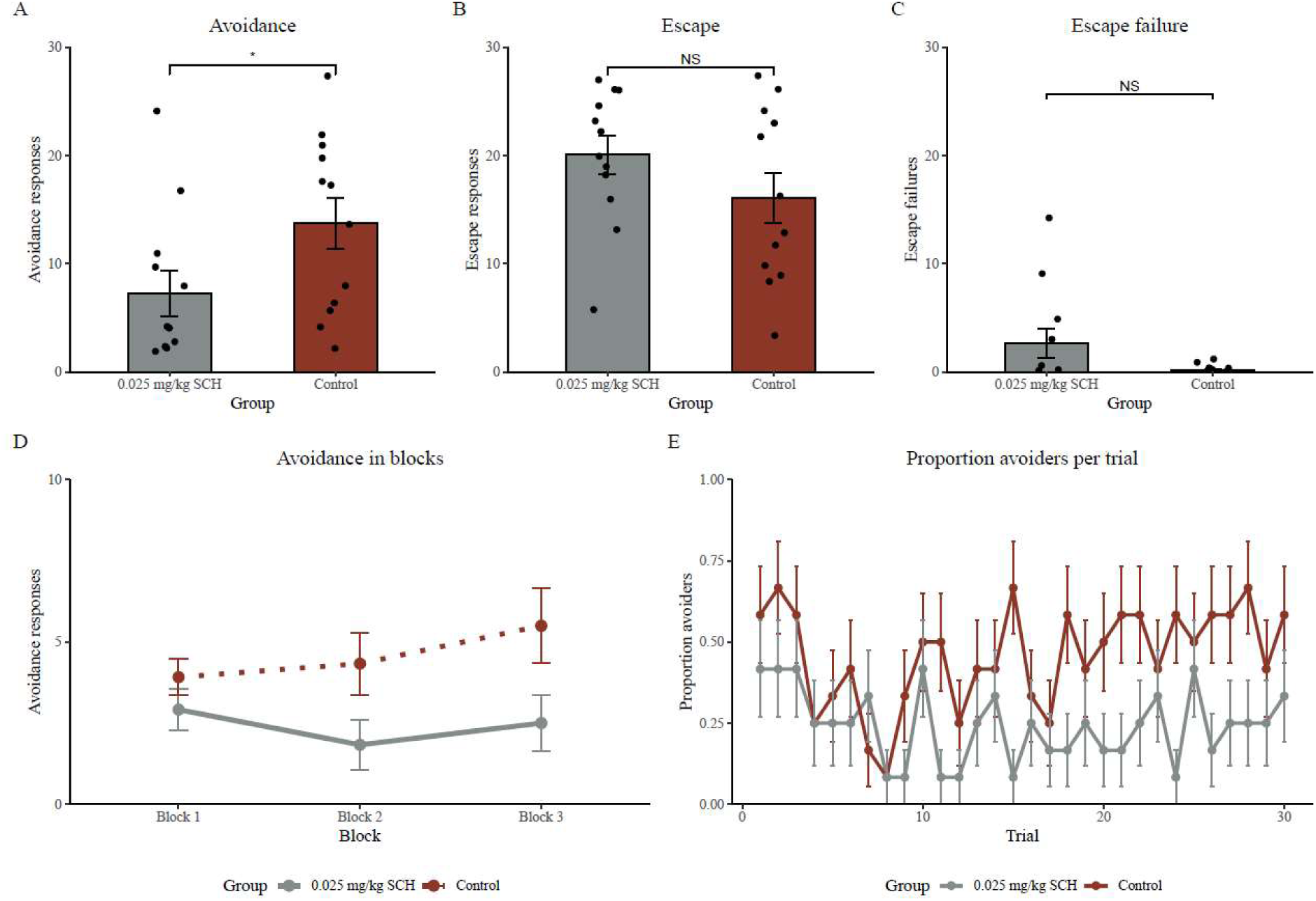
Two-way active avoidance training of Experiment 2. (A) Mean (± SEM) number of avoidance responses, (B) escape responses and (C) escape failures for the 0.025 mg/kg SCH (n = 12) and control (n = 12) groups. * *p* < .05 (D) Mean (± SEM) number of avoidance responses in blocks of 10 trials for the 0.025 mg/kg SCH and control groups. (E) Proportion of avoiders per trial ± SEM for the 0.025 mg/kg SCH and control groups.

When analyzing the data of the avoidance training session in three blocks of 10 trials each, there was a trend towards a main effect of Group (*F*(1,22) = 4.217, *p* = .052, ƞ*_p_²* = 0.161), but no significant main effect of Block (*F*(1.41,31.09) = 1.785, *p* = .191, ƞ*_p_²* = 0.075), nor a significant Group by Block interaction effect (*F*(1.41,31.09) == 2.246, *p* = .136, ƞ*_p_²* = 0.093; Figure 4D). Finally, we conducted an exploratory generalized linear mixed model (GLMM) analysis to examine the effects of Trial and Group (reference group: control group) on the binary avoidance data (avoidance = 1, escape/escape failure = 0), with a random intercept for each subject. The logistic regression coefficients indicated a significant effect of Trial (*β* = 0.033, *SE* = 0.014, *Z* = 2.320, *p* = .020), indicating that as the trials progressed, the likelihood of performing an avoidance response increased. Furthermore, although the main effect of Group was not statistically significant (*β* = -0.527, *SE* = 0.668, *Z* = -0.789, *p* = .430), the interaction between Trial and Group SCH 23390 was significant (*β* = -0.049, *SE* = 0.022, *Z* = -2.254, *p* = .024), suggesting that while the number of avoidance responses increased with trials, this effect was smaller in the SCH 23390 group. Together, these findings suggest that 0.025 mg/kg was effective in reducing the number of avoidance responses performed during acquisition, in line with our hypothesis.

#### 3.2.2. Two-way active avoidance test

In contrast to the acute SCH effect seen during avoidance acquisition, the drug-free avoidance test 24 hours later showed no significant difference in the number of avoidance responses between the SCH and vehicle groups (*t*(22) = -0.912, *p* = .814, *d* = -0.372; Figure 5A). Furthermore, there was no difference in the number of escape responses (*t*(22) = 0.792, *p* = .437, *d* = 0.414; Figure 5B), nor in the number of escape failures (*t*(22) = 0.993, *p* = .332, *d* = 0.405; Figure 5C) during the avoidance test.

**Figure 5.**
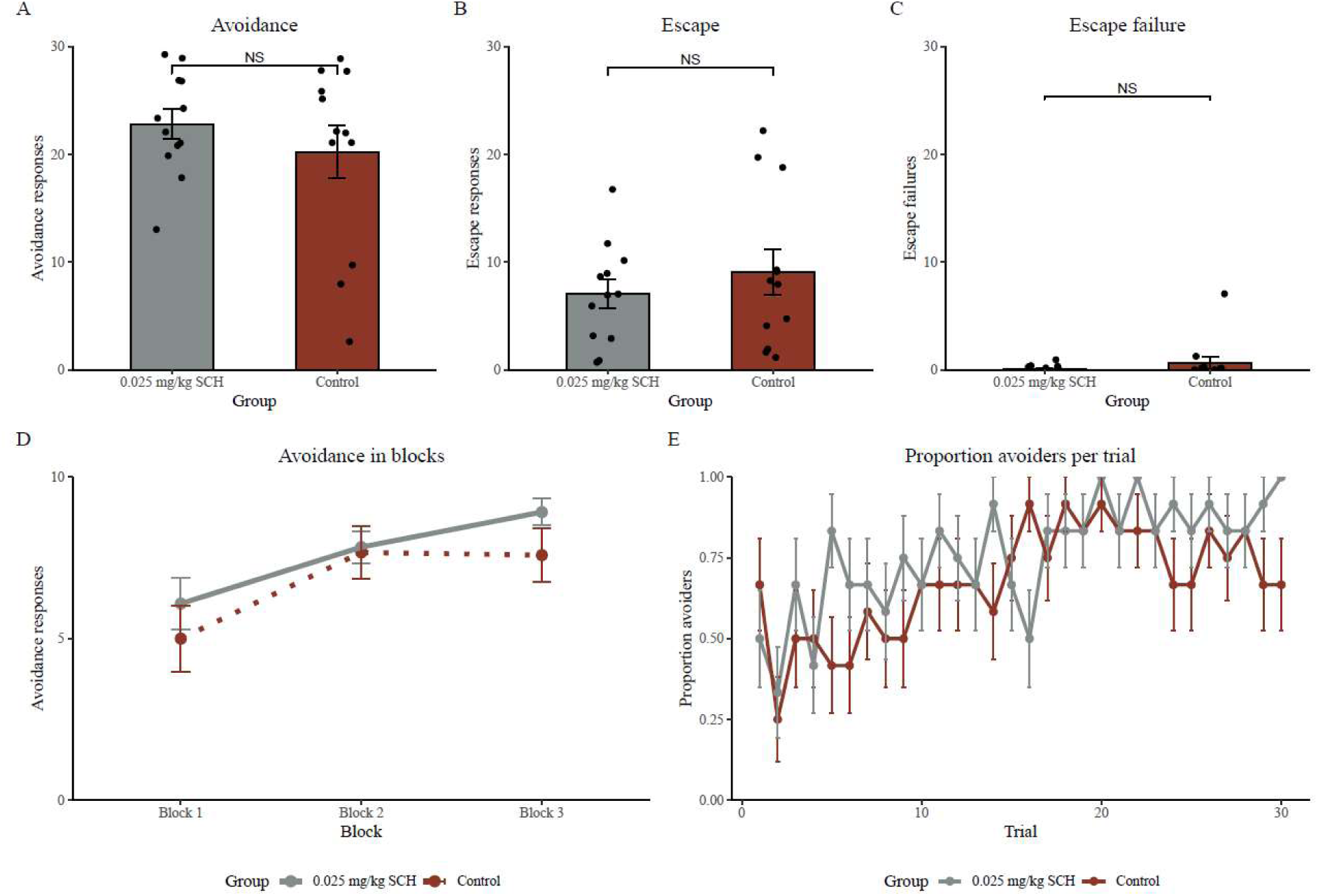
Two-way active avoidance test of Experiment 2. (A) Mean (± SEM) number of avoidance responses, (B) escape responses and (C) escape failures for the 0.025 mg/kg SCH (n = 12) and control (n = 12) groups. (D) Mean (± SEM) number of avoidance responses in blocks of 10 trials for the 0.025 mg/kg SCH and control groups. (E) Proportion of avoiders per trial ± SEM for the 0.025 mg/kg SCH and control groups.

Similarly, when analyzing the data of the avoidance test in three blocks of 10 trials each, there was no significant main effect of Group (*F*(1,22) = 0.831, *p* = .372, ƞ*_p_²* = 0.036), only a significant main effect of Block (*F*(2,44) = 22.431, *p* < .001, ƞ*_p_²* = 0.505; Figure 5D). These findings suggest that the rats performed the avoidance response equally well in both groups. Moreover, an exploratory generalized linear mixed model (GLMM) analysis was applied to examine the effects of Trial and Group (reference group: control group) on the binary avoidance data (avoidance = 1, escape/escape failure = 0), with a random intercept for each subject. The logistic regression coefficients indicated a significant effect of Trial (*β* = 0.090, *SE* = 0.018, *Z* = 5.101, *p* < .001), suggesting that avoidance still increased across trials. The main effect of Group (*β* = 0.357, *SE* = 0.671, *Z* = 0.532, *p* = .595), as well as the Group by Trial interaction (*β* = 0.010, *SE* = 0.025, *Z* = 0.422, *p* = .673) were not significant. These results indicate a consistent positive association between Trial and the likelihood of an avoidance response, independent of the group. Finally, an explorative analysis on the first three trials of the test session indicated no significant difference between the SCH group and the control group (*t*(22) = -0.205, *p* = .840, *d* = -0.083), suggesting that, even early on in the test session, the performance of both groups was similar. Moreover, when evaluating the number of shuttles during the ITIs throughout the test session, we did not observe a significant difference in shuttles between the 0.025 mg/kg SCH group and the control group (*t*(22) = 0.677, *p* = .505, *d* = 0.276).

Overall, these findings suggest that, although SCH significantly reduces avoidance responding during avoidance acquisition, this effect did not persist during a drug-free test under continued reinforcement on the next day. These results seem inconsistent with an impairment in avoidance learning through D1R antagonism, as the performance of SCH rats did not differ from that of the control group at the start nor throughout the drug-free test. To further corroborate these results, we performed an exploratory mixed ANOVA with repeated measures factor Phase (Block 3 of avoidance training session (last 10 trials) and Block 1 of avoidance test (first 10 trials) and between-subjects factor Group (0.025 mg/kg SCH and control). We observed a significant Phase by Group interaction effect (*F*(1,22) = 6.184, *p* = .021, ƞ*_p_²* = 0.219). Post-hoc testing showed that rats of the SCH group displayed a significant increase in avoidance responses from the end of avoidance training to the beginning of the avoidance test (*t* = -3.086, *p* = .032), whereas the control group did not show this increase from training to test (*t*( = 0.431, *p* = 1). Again, these findings seem inconsistent with an impairment of avoidance learning proper through D1R blockade, but rather point towards an acute impairment of secondary processes during the training session.

#### 3.2.3. Locomotor activity and rotarod test

To evaluate whether there were any acute effects of 0.025 mg/kg SCH 23390 on locomotor activity, we first evaluated the number of crossings during the ITIs of avoidance acquisition. We did not observe a significant difference in the number of crossings between the SCH group and the control group (*t*(22) = 1.881, *p* = .073, *d* = 0.768, Figure 6A), but the data did indicate a trend towards fewer crossings for the SCH group. To further evaluate this potential motor effect, all rats were subjected to an accelerating rotarod that took place 7 days after avoidance training. Additionally, a group of rats that received 0.05 mg/kg SCH was also included to further validate the findings observed in the locomotor activity test of Experiment 1. In contrast to a general locomotor activity test in an open arena, which also measures exploratory behavior, the accelerating rotarod is a more specific test of motor function.

**Figure 6.**
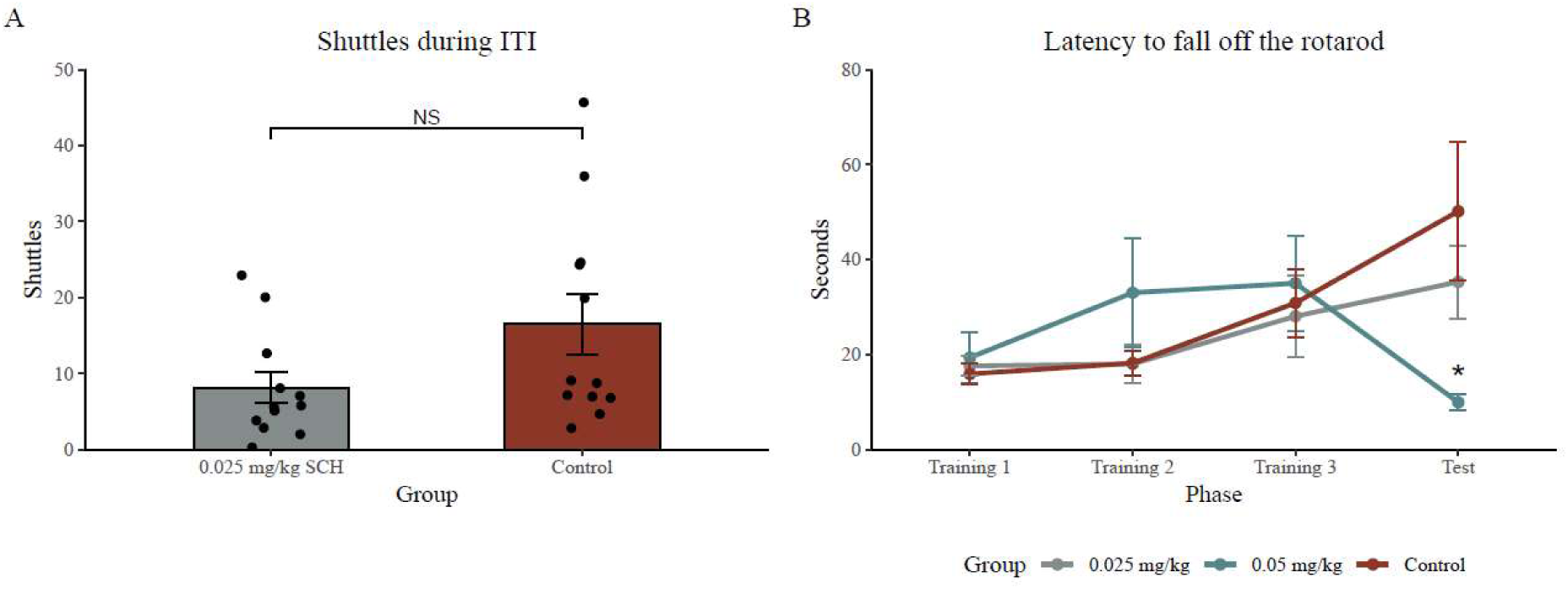
(A) Mean number of shuttles ± SEM during the ITIs of the avoidance training session for the 0.025 mg/kg SCH (n = 12) and control (n = 12) groups. (B) The mean number of seconds the rats remained on the accelerating rotarod ± SEM for the three days of training and the test day for the 0.05 mg/kg SCH (n = 8), 0.025 mg/kg SCH (n = 8) and control (n = 8) groups. * *p* < .05.

All rats first underwent three days of rotarod training. On the fourth day, the rats were administered with either 0.05 mg/kg SCH 23390 (n = 8), 0.025 mg/kg SCH 23390 (n = 8), or vehicle (0.9% physiological saline + 17% DMSO, n = 8) 20 minutes prior to the rotarod test. Throughout the three days of rotarod training, performances increased for all groups similarly, as illustrated by a significant main effect of Training session (*F*(2,42) = 7.091, *p* = .002, ƞ*_p_²* = 0.252), and a non-significant main effect of Group (*F*(2,21) = 0.597, *p* = .560, ƞ*_p_²* = 0.054; Figure 6B). These findings suggest that all groups acquired a similar rotarod performance.

When comparing the final day of rotarod training with the test day, there was a significant Group by Phase interaction effect (*F*(2,21) = 4.194, *p* = .029, ƞ*_p_²* = 0.285; Figure 6B). Post-hoc testing illustrated a significant decrease in performance for rats that received 0.05 mg/kg SCH, compared to the vehicle group (*p* = .038). Contrarily, 0.025 mg/kg SCH did not impair performance compared to the vehicle group (*p* = .858). These findings suggest that while 0.05 mg/kg SCH significantly impaired motor function, 0.025 mg/kg SCH did not.

## 4. Discussion

The current study was set up to test the involvement of D1R and D2R neurotransmission in the acquisition of two-way active avoidance in rats. Previous studies observed a reduction in avoidance responses when D1R or D2R antagonists were administered before avoidance training (Reis et al., 2004; Boschen et al., 2011; Wietzikoski et al., 2012). Importantly, although these findings were taken as evidence that DR blockade directly impairs avoidance acquisition, such an impairment in learning should also decrease performance on a later drug-free test, at least initially, a notion that has largely gone untested, and when tested, has yielded diverging results. Therefore, while Experiment 1 served to replicate the acute effects of D1R and D2R blockade on the acquisition of avoidance (Wietzikoski et al., 2012; Reis et al., 2004; Carvalho et al., 2009), Experiment 2 included a drug-free follow-up test under continued reinforcement. Our findings showed that, although D1R blockade impaired the performance of avoidance during avoidance training, this impairment did not persist the next day, thus suggesting that D1R blockade did not impair instrumental learning per se.

Experiment 1 aimed to replicate the finding that blocking D1 or D2 receptors prior to avoidance training acutely impairs the development of avoidance (Reis et al., 2004; Wietzikoski et al., 2012). Our results indeed showed fewer avoidance responses following administration of D1R antagonist SCH 23390 (0.05 mg/kg), compared to a vehicle group. Further exploration of the data revealed that, in addition to a reduction in successful CS-elicited avoidance responses, US-elicited escape responses were also less frequent following SCH administration. This was further paralleled by an overall reduction in movement, which invites an alternative explanation for the reduction of avoidance responses in terms of impaired motivation or locomotor ability. Motivation and locomotor abilities are both closely tied to instrumental learning. If rats cannot move effectively, or do not want to, they might not be able to acquire the relevant associations between the warning signal and the behavioral response that is required to prevent the aversive outcome.

In contrast to the reduction in avoidance responses that we observed after D1R blockade, D2R antagonist sulpiride (20 mg/kg) did not significantly affect the number of avoidance responses during training, compared to a vehicle group. The same dose of sulpiride did impair avoidance in previous studies (Carvalho et al., 2009; Reis et al., 2004), but there are many differences in experimental parameters that could explain the divergent effects, including the number of trials, the intertrial interval duration, the nature of the CS, no overlap between the CS and US, and the presentation of a brief safety signal after successful avoidance or escape responses. D2 receptor blockade may also have less of an effect on avoidance acquisition due to the differential role of D1R and D2R in evaluating positive versus negative situations. D1Rs are mainly expressed on direct pathway medium spiny neurons, whereas D2Rs are mainly expressed on indirect pathway neurons, both of which project from the striatum to the basal ganglia thereby contributing to the facilitation or suppression of body movements, respectively (Kravitz et al., 2012). When a behavioral response is followed by an outcome that was better than expected (reward PE), dopamine neurons will fire a burst of spikes resulting in the activation of D1Rs on the direct pathway, thereby reinforcing cortico-striatal synapses to promote instrumental learning (Bromberg-Martin et al., 2010). D2Rs, on the other hand, are mostly involved in the suppression of actions when an outcome was worse than expected. Given the notion that the omission of an expected aversive event is an outcome that was better than expected (rPE; Kim et al., 2006), it is reasonable to assume that D1R are more prominently involved in this process than D2R.

In Experiment 2, a lower dose of the D1R antagonist SCH 23390 (0.025 mg/kg) that did not significantly affect locomotor activity in an accelerating rotarod test, was used to further evaluate the effect of D1R blockade on avoidance learning. In prior work, this dose has been found to acutely impair the acquisition of avoidance (Diaz-Veliz et al., 1998; Reis et al., 2004), but none of these studies included a follow-up test under continued reinforcement to support the conclusion that SCH 23390 impairs instrumental learning proper. While our results confirmed a reduction in acquisition of avoidance following 0.025 mg/kg SCH 23390 administration, this reduction was not maintained on a drug-free test 24 h later, whether evaluated across the entire test session or specifically during the early trials only. These findings suggest that, rather than impairing avoidance acquisition directly, SCH 23390 might have acutely affected secondary processes during avoidance learning. Besides motor function, which was not significantly impaired during the accelerated rotarod test by 0.025 mg/kg SCH administration, dopamine is also critically involved in motivation. Previous studies observed that rats with lesions in the striatal dopamine system (caused by 6-OHDA) still consume food and even enjoy it (Berridge et al., 1989), but they will not exert effort to actively obtain rewards (Berke, 2018). In the context of avoidance, D1R blockade could have caused a reduction in the motivation to actively avoid foot shock, as rats need to exert some effort to shuttle to the opposite compartment. Indeed, previous studies observed a decrease in food-or water-rewarded lever pressing following SCH 23390 administration (Ljungberg, 1988; Beninger et al., 1987), thereby highlighting a potential role for D1 receptors in motivational processes. Such disruption in motivation, perhaps combined with a subtle reduction in locomotor activity (Suppl. Fig. 2B, 3A), could acutely result in less avoidance responses, even when the learning is intact. If learning per se is indeed not impaired, this would result in intact performance during a drug-free test, which is what we observed in Experiment 2.

The conclusion that the effects of D1R blockade reflect a motivational deficit rather than a learning deficit needs to be made with caution, because this was not the research question at hand and requires further investigation. Furthermore, it should be noted that the control group showed limited progress in avoidance learning during training in Experiment 2, as there was no significant main effect of Block (Fig. 4D, dotted line). This suggests that the control group of Experiment 2 did not fully acquire the avoidance response during avoidance training and continued to learn during the drug-free reinforced test (Fig. 5D). The relatively low number of avoidance responses in the control group may theoretically reduce the possibility of observing significant differences in performance between the control rats and the drug-treated rats. The significant group difference during training (Fig. 4A) does, however, argue against such an issue. Even with the relatively low levels of responding in the control group, we still had enough power to show a significant deficit in the SCH rats. Moreover, we also observed a significant interaction when comparing avoidance responses at the end of training, with the avoidance responses at the start of the test. Rats that were administered with 0.025 mg/kg SCH increased their avoidance responding from training to test, whereas the control group did not, thereby further supporting our notion that the D1R blockade did not affect instrumental learning proper.

In summary, our findings suggest that systemic administration of 0.025 mg/kg SCH 23390 reduces the number of avoidance responses during learning, but may not impair avoidance acquisition per se, in contrast to what previous literature has put forward. Our data show that the rats’ performance was similar to the performance of a vehicle group during a drug-free avoidance test, despite the lower performance of SCH rats observed during training. We propose that effects of D1R blockade might reflect an acute impairment in dopamine-mediated motivation, rather than an effect on avoidance learning directly.

## CRediT Authorship contribution statement

Laura Vercammen: conceptualization, data curation, formal analysis, funding acquisition, investigation, methodology, visualization, writhing – original draft, writing – review and editing; Alba Lopez-Moraga: formal analysis, visualization, writing – review and editing; Tom Beckers: conceptualization, methodology, supervision, writing – review and editing; Bram Vervliet: conceptualization, methodology, supervision, writing – review and editing, funding acquisition; Laura Luyten: conceptualization, methodology, supervision, writing – review and editing

