## Supplementary Material for "The impact of systemic blockade of dopamine receptors on the acquisition of two-way active avoidance in rats"

The supplementary material contains all preregistered analyses that were not included in the manuscript. The average latency to avoid the US, as well as the average latency to escape the US were analyzed using two-sided independent samples t-tests (Suppl. Fig. 1). Moreover, the latency to avoid the US was calculated per block of 10 trials within a session and analyzed using a mixed design ANOVA (repeated-measures factor: Block, between-subjects factor: Group). The average number of seconds of freezing during CS1 and during each even CS presentation (CS2, CS4,…) was analyzed using a mixed design ANOVA (repeated-measures factor: CS, between-subjects factor: Group). Moreover, the average number of seconds of freezing across all even CS presentations was analyzed using two-sided independent samples t-tests (Suppl. Fig. 2). Finally, the average number of shuttles during the 5-minute habituation of both the avoidance training session and the avoidance test session of Experiment 2 were analyzed using two-sided independent samples t-tests (Suppl. Fig. 3).

### 1. Avoidance and US latencies

#### 1.1. Average latency to avoid the US and the average latency to escape the US

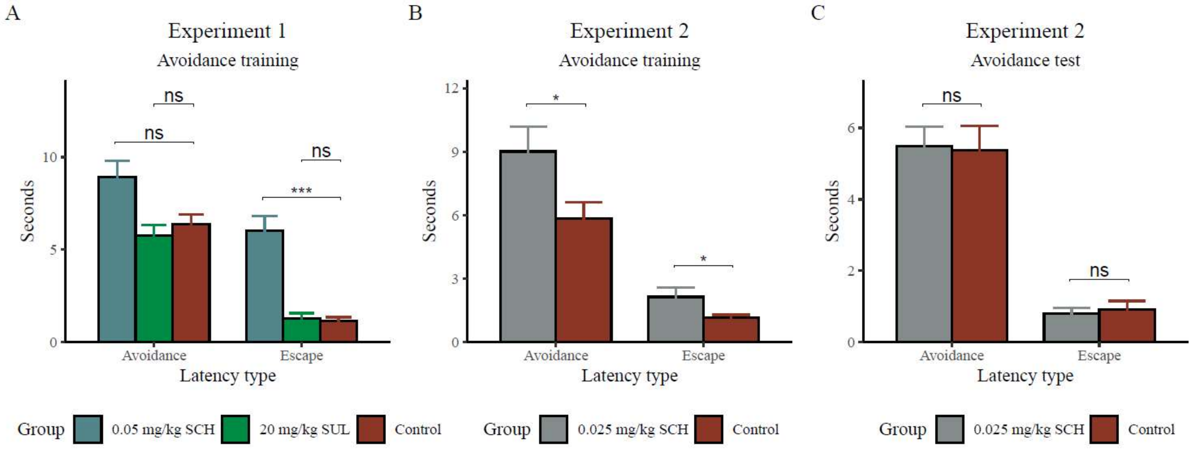


*Supplementary Figure 1*. Mean latency (± SEM) to avoid the aversive stimulus (avoidance) and to escape the aversive stimulus (escape) for (A) the avoidance training session of Experiment 1, (B) the avoidance training session of Experiment 2, and (C) the avoidance test session of Experiment 2. Note that the CS had a maximum duration of 20 s and the US a maximum duration of 10 s. * $p < .05$, *** $p < .001$.

Experiment 1 avoidance training (Suppl. Fig. 1A). There was a significant difference in both the latency to avoid ($t(21) = -2.452$, $p = .023$, $d = -1.024$) the foot shock and the latency to escape ($t(21) = -6.269$, $p < .001$, $d = -2.617$) between rats of the 0.05 mg/kg SCH group (n = 12) and the control group (n = 12). There were no significant differences in the latency to avoid ($t(22) = 0.785$, $p = .441$, $d =$

0.320), nor in the latency to escape ($t(22) = -0.364$, $p = .719$, $d = -0.149$) for rats of the 20 mg/kg SUL group (n = 12) compared to the control group (n = 12).

Experiment 2 avoidance training (Suppl. Fig. 1B). There was a significant difference in the latency to avoid the foot shock ($t(21) = -2.307$, $p = .031$, $d = 0.963$) and the latency to escape the foot shock ($t(21) = 2.272$, $p = .034$, $d = 0.948$) for rats of the 0.025 mg/kg SCH group (n = 12), compared to the control group (n = 12).

Experiment 2 avoidance test (Suppl. Fig. 1C). There were no significant differences in the latency to avoid the foot shock ($t(22) = 0.143$, $p = .887$, $d = 0.059$) nor in the latency to escape the foot shock ($t(22) = -0.368$, $p = .716$, $d = -0.150$) for rats of the 0.025 mg/kg SCH group (n = 12), compared to the control group (n = 12).

**1.2. Average latency to avoid the US and average latency to escape the US per block of 10 trials**

Experiment 2 avoidance training. The preregistered analysis on the latencies per block of 10 trials was not conducted for the avoidance training session of Experiment 2 due to the exclusion of multiple subjects that failed to avoid within a block (42 %), resulting in NA values for the avoidance latency.

Experiment 2 avoidance test. The preregistered analysis on the latencies per block of 10 trials was conducted for the avoidance test of Experiment 2. Two subjects were excluded from the analysis due to a failure to avoid within a block, resulting in NA values for the avoidance latency. The results showed no significant effect of Block ($F(1.52,30.41) = 2.353$, $p = .124$, $\eta_p^2 = 0.105$), no significant main effect of Group ($F(1,20) = 0.569$, $p = 0.460$, $\eta_p^2 = 0.028$), and no significant Group by Block interaction effect ($F(1.52,30.41) = 0.797$, $p = 0.428$, $\eta_p^2 = 0.038$).

### 2. Freezing

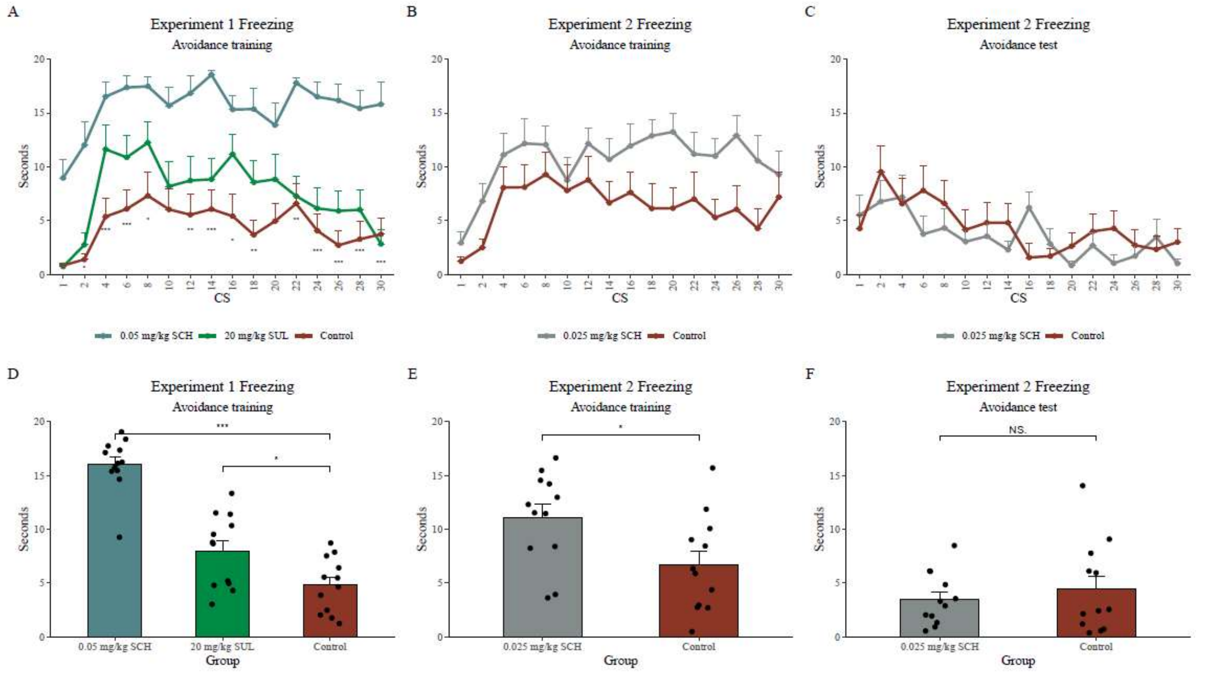


*Supplementary Figure 2.* Average number of seconds of freezing during CS1 and during each even CS presentation (CS2, CS4,…) ± SEM for each group separately during (A) avoidance training of Experiment 1, (B) avoidance training of Experiment 2, and (C) the avoidance test of Experiment 2. Average number of seconds of freezing across all even CS presentations ± SEM for each group separately during (D) avoidance training of Experiment 1, (E) avoidance training of Experiment 2, and (F) the avoidance test of Experiment 2. *** $p < .001$, * $p < .05$.

Experiment 1 avoidance training. When evaluating freezing throughout the session, we observed a significant main effect of CS ($F(9.15,301.82) = 6.648$, $p < .001$, $\eta_p{}^2 = 0.168$, and a significant main effect of Group ($F(2,33) = 51.959$, $p < .001$, $\eta_p{}^2 = 0.759$), but no significant Group by CS interaction effect ($F(18.29,301.82) = 0.859$, $p = .631$, $\eta_p{}^2 = 0.049$; Suppl. Fig. 2A). Post-hoc testing showed significant differences between the 0.05 mg/kg SCH group (n = 12) and the control group (n = 12) for all CS presentations, except during CS1, CS10 and CS20. Moreover, on average, post-hoc testing showed a significant difference between the 0.05 mg/kg SCH group and the control group ($t = 9.853$, $p < .001$), as well as between the 20 mg/kg SUL group (n = 12) and the control group ($t(2.661$, $p = .012$). When the number of seconds of freezing was averaged across all even CS presentations, we observed a significant difference between the 0.05 mg/kg SCH and control group ($t(21.994) = 10.869$, $p < .001$, $d = 4.437$), as well as between the 20 mg/kg SUL and the control group ($t(22) = 2.597$, $p = .016$, $d = -1.060$; Suppl. Fig. 2D), suggesting that, even though we did not observe a significant difference in avoidance responding between the SUL and control group, there is a significant effect on freezing, with the SUL group freezing more than the control group.

Experiment 2 avoidance training. When evaluating freezing throughout the session, we observed a significant main effect of CS ($F(8.1,178.2) = 4.104$, $p < .001$, $\eta_p^2 = 0.157$) and a significant main effect of Group ($F(1,22) = 6.299$, $p = .02$, $\eta_p^2 = 0.223$), but no significant Group by CS interaction effect ($F(8.1,178.2) = 0.733$, $p = .664$, $\eta_p^2 = 0.032$; Suppl. Fig. 2B). Post hoc testing showed a significant overall effect of Group ($t = -2.510$, $p = 0.020$), but comparisons for each CS separately were not significant when corrected for the number of tests that were run. Moreover, when averaging the number of seconds of freezing across all even CS presentations, we observed significantly more freezing for the 0.025 mg/kg SCH group (n = 12), compared to the control group (n = 12; $t(22) = 2.47$, $p = 0.022$, $d = -1.010$; Suppl. Fig. 2E).

Experiment 2 avoidance test. When evaluating freezing throughout the test session, we observed a significant main effect of CS ($F(6.26,137.77) = 4.468$, $p < .001$, $\eta_p^2 = 0.169$), but no significant main effect of Group ($F(1,22) = 0.413$, $p = .527$, $\eta_p^2 = 0.018$), nor a significant Group by CS interaction effect ($F(6.26,137.77) = 1.580$, $p = .154$, $\eta_p^2 = 0.067$; Suppl. Fig. 2C). Moreover, when the number of seconds of freezing was averaged across all even CS presentations (including C1, given that rats have already experienced the CS-US association during training on the previous day), we observed no significant difference in freezing between the 0.025 mg/kg SCH group (n = 12) and the control group (n = 12; $t(17.53) = -0.642$, $p = .529$, $d = 0.2622$).

The interpretation of the freezing values is challenging, because they are analyzed as seconds of freezing and not as percentages (of the varying CS durations), but this method has been used previously in the literature (e.g., Moscarello & Ledoux, 2013).

### 3. Shuttles during Habituation

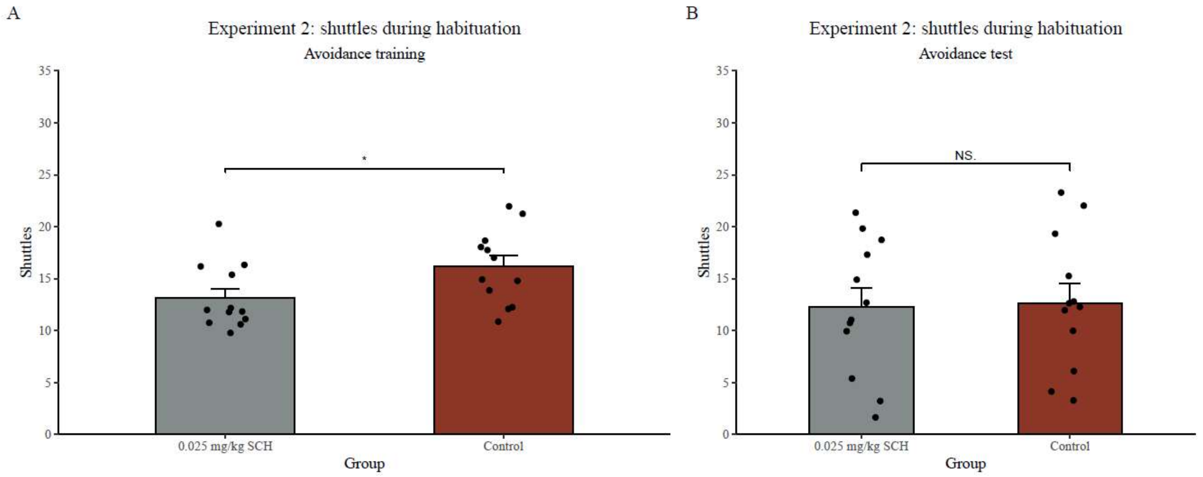


*Supplemental Figure 3.* Average number of shuttles (± SEM) during the first 5 minutes of habituation for (A) the avoidance training session and (B) the avoidance test session of Experiment 2. * $p < .05$.

When evaluating the number of shuttles during the first five minutes of the avoidance training session of Experiment 2, we observed a significant difference, with the 0.025 mg/kg SCH group shuttling significantly less compared to the control group ($t(22) = 2.237$, $p = .036$, $d = 0.913$; Suppl. Fig. 3A). In contrast, during the first five minutes of habituation of the avoidance test of Experiment 2, there was no significant difference in the number of shuttles between the 0.025 mg/kg SCH group and the control group ($t(22) = 0.677$, $p = .505$, $d = 0.276$; Suppl. Fig. 3B).

Together, the small effects on freezing (Suppl. Fig. 2B) and the shuttles during the 5-minute habituation period (Suppl. Fig. 3A) are in line with the notion that there might be a combination of a (subtle) motor effect and an effect of motivation during avoidance training, instead of a pure motivation effect. At the same time, it is, of course, difficult to disentangle them, even more in the avoidance context - although the group difference already seems to be there before shocks are given (in the shuttles and on CS1).
